# Rapid expression of pyruvate decarboxylase from *Zymomonas mobilis* in *E. coli* BL21 LysY/I^q^

**DOI:** 10.1101/2025.05.25.654064

**Authors:** S. Bilal Jilani, Daniel G. Olson

## Abstract

*E. coli* BL21 DE3 strain is commonly used to express and purify non-toxic prokaryotic proteins in high yields. Traditionally, IPTG based induction of the host strain at reduced temperatures for extended period (12-16 h) is performed to obtain high yield of functional proteins. It is desirable to explore methods which result in high yield of protein within a short period of time. We report rapid purification of pyruvate decarboxylase (PDC) enzyme from *Zymomonas mobilis* using *E. coli* BL21 pLysY/I^q^ as a host strain. High yield of purified PDC at 0.33 (±0.02) mg was obtained after two-hour induction by 0.6 mM IPTG at 37^°^C. The enzyme yield was comparable to 0.37 (±0.08) mg obtained in *E. coli* BL21 DE3 strain (used as control) after 16 h induction by 0.6 mM IPTG at 18^°^C. Similar values of the maximum specific activity of the enzyme expressed and purified at 37^°^C and 18^°^C were obtained at 78.31 (±1.13) in strain LysY/I^q^ and 85.73 (±4.39) µmol/min/mg protein in strain DE3, respectively. In almost all IPTG treatments, the kinetic parameters of the purified enzyme - *app K*_*m*_, *app V*_*max*_, *K*_*cat*_ and *K*_*cat*_*/K*_*m*_ -also did not vary remarkably between the two temperature regimes. Based upon the data presented here, we propose that *E. coli* BL21 LysY/I^q^ strain has potential to serve as a host for efficient and rapid expression (2 h) of non-toxic proteins. Results of this study will aid in cell free system study which require rapid scale up of the complexity of metabolic pathways by utilizing multiple purified enzymes involved in bioconversion of the substrate of interest.

## Introduction

Cell free biology (Jewett et al., 2008) has opened an important avenue for investigation of complex metabolic pathways of non-model organisms which are not readily amenable to genetic manipulations (Chubukov et al., 2018; Cui et al., 2020; Jilani et al., 2024). Cell free experimental investigations cfrequently involve heterologous purification of enzymes which are required for perturbing the cell free system and to evaluate the fate of targeted substrate. Purification of proteins is a cumbersome procedure and is a major hindrance in scaling up the complexity of multistep metabolic pathways, under *invitro* conditions, involving purification of several enzymes (Bowie et al., 2020; Chae et al., 2017; Guterl et al., 2012). Extended cultivation of host strains at reduced temperatures is commonly performed to prevent degradation of the polypeptide and promote proper folding to obtain high yields of the functional heterologous protein (Farewell & Neidhardt, 1998; Peti & Page, 2007; Schein & Noteborn, 1988; Vera et al., 2007). It is desirable to explore strategies where a heterologous protein can be rapidly expressed without compromising on its yield or activity, which will in turn contribute towards reducing the experimental turnaround time.

*E. coli* BL21 DE3 strain is commonly used for obtaining high yields of heterologous protein (Rosano & Ceccarelli, 2014). It harbors lambda DE3 prophage which carries the gene for T7 RNA polymerase (Studier & Moffattf, 1986). The expression of the polymerase is controlled by a lacUV5 promoter which can be induced by IPTG. It allows high level expression of the polymerase which in turn leads to transcription of the cloned gene sequence whose promoter can be recognized by the T7 RNA polymerase. Hereafter referred to as DE3. On the other end of the expression levels, *E. coli* BL21 LysY/I^q^ strain lacks a lambda prophage and harbors a mutation in the promoter region of lacI gene which allows high expression levels of the lacI repressor which binds to the DNA operator sites and prevent initiation of transcription by T7 RNA polymerase in absence of IPTG. In presence of IPTG, expression of Lys Y lysozyme is induced which leads to degradation of lacI and allows T7 RNA polymerase to bind to the operator sites and initiate transcription of the cloned gene of interest. Hereafter referred to as LysY/I^q^. Commonly, strain LysY/I^q^ is used for expression and purification of proteins which have toxic effect on microbial physiology and result in relatively lower yields of the heterologous protein.

Pyruvate dehydrogenase (PDC) from *Zymomonas mobilis* is an important protein from industrial perspective (Conway et al., 1987; Hoppner & Doelle, 1983; Neale et al., 1987). PDC catalyzes non-oxidative decarboxylation of pyruvate to yield acetaldehyde which is subsequently converted to ethanol to yield NAD^+^ (Fig 1). Ethanol is subsequently used as a blending agent in gasoline. Multiple studies have been carried out where PDC has been expressed *invivo* in *E. coli* strains of industrial utility (Alterthum’ And & Ingram2, 1989; Ingram,’ et al., 1987; Lindsay et al., 1995; Ohta et al., 1991) to produce bioethanol. There are no published reports of the PDC being toxic to the microbial cell. It is a stable protein which can withstand elevated temperatures of 60^°^C without losing any activity (Raj et al., 2002). These characteristics make PDC an ideal candidate in our studies which reduces probability of confounding factors from influencing our observations. In the reported literature the purification of PDC has been performed after extended induction time period and there is lack of clarity on the yield of the purified protein.

**Figure 1:**
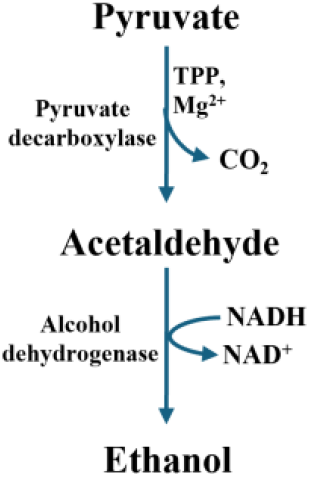
Reaction catalyzed by pyruvate decarboxylase (PDC). Activity of PDC requires presence of thiamine pyrophosphate (TPP) and Mg^2+^. Acetaldehyde is further reduced to ethanol by alcohol dehydrogenase coupled with oxidation of NADH to NAD^+^.

In this work, we investigated the optimum cultivation conditions and host strain which results in highest yields of functional PDC protein. Based on our results, we report two main observations. Firstly, two hour cultivation of *E. coli* host strains at 37^°^C result in comparable yield and specific activity of PDC as that obtained after 16 h cultivation at 18^°^C. Secondly, at 37^°^C, *E. coli* BL21 strain LysY/I^q^ result in highest yield and specific activity of PDC. While at 18^°^C, E. coli strain DE3 results in highest yield and specific activity of purified PDC.

## Materials and Methods

### Strains and cultivation conditions

Protein expression was tested in two microbial strains of *E. coli* BL21 DE3 with genotype F^-^ *omp*T *hsd*SB (rB-mB-) *gal dcm* (DE3) and LysY/I^q^ strain deficient in λ-prophage. High copy number plasmid pET22b harboring pyruvate decarboxylase (PDC) sequence derived from *Zymomonas mobilis* was used for PDC expression and purification. pET22b-PDC was transformed in respective strain by chemical transformation and one well isolated colony each of *E. coli* BL21 DE3 – PDC and *E. coli* BL21 LysY/I^q^ – PDC was used for protein expression and purification. All cultivation was performed in LB media in presence of 100 µg/mL ampicillin at either 37^°^C or at 18^°^C with shaking at 200 RPM.

### PDC expression

Primary cultures of BL21 DE3 – PDC and BL21 LysY/I^q^ – PDC strains were grown separately overnight at 37^°^C. 1% of the saturated overnight culture of each strain was used to inoculate six tubes (50 mL capacity) containing 5 mL of LB media. The cultures were allowed to grow at 37^°^C until OD_600_ reached 0.4. Then tubes of respective strain was induced with either 0, 0.1, 0.3, 0.6, 1.2 or 2.4 mM IPTG. One set of each strain was induced for 2 h at 37^°^C and another set was induced for 16 h at 18^°^C. The cells were harvested after induction period by pelleting at 4,400 RPM at 4^°^C and stored at -80^°^C until further analysis. All SDS-PAGE analysis was performed in gels with 10-16% acrylamide content.

### PDC purification

The cell pellets were thawed on ice and resuspended in lysis buffer consisting of 5 mM imidazole, 500 mM NaCl, 20 mM Tris-HCl (pH 7.2), 1 mg/mL lysozyme and 1 mM phenylmethylsulfonyl fluoride. The cells were lysed using a microtip sonicator (Misonix S-4000, 600W maximum output power) at 30% amplitude with a 6 s on/off cycle for 1 min. The lysate was centrifuged at 4 ^°^C for 30 minutes at 14,100 X g. The supernatant was bound with nickel-charged affinity resin (Ni-NTA) at 4 ^°^C on a nutator mixer for around 12 h. The bound protein was passed through a gravity column (Bio-Rad) and the resin was washed with a buffer containing 15 mM imidazole, 500mM NaCl and 20 mM Tris-HCl (pH 7.2). Bound protein was eluted with a buffer containing 650 mM imidazole, 500 mM NaCl, and 20 mM Tris-HCl (pH 7.2). The eluted protein was dialyzed against 100 mM phosphate buffer (pH 6.0). Purity of the protein was confirmed by SDS-PAGE (10-16% acrylamide) analysis. Protein concentration was estimated by bicinchoninic acid (BCA) assay (G-Biosciences) using bovine serum albumin as standard.

### Specific activity of PDC

Specific activity in the cell lysates was determined via a coupled assay with Adh enzyme in HP-Agilent (Model 8453) UV-Vis diode array spectrophotometer. Assay was performed in a 1 mL reaction mixture consisting of 50 mM phosphate buffer (pH 6.0), 20 mM sodium pyruvate, 1 mM MgCl2, 1 mM thiamine pyrophosphate, 5 mM NADH and 75 U of Adh. The reaction constituents were added in the following sequence – to the phosphate buffer, MgCl2 was added, followed by thiamine pyrophosphate, sodium pyruvate, NADH and Adh. Then absorbance at 390 nm was allowed to stabilize and finally reaction initiated by addition of either cell lysate or purified PDC and decrease of 390 nm signal corresponding to disappearance of NADH was monitored. In order to inactivate non specific enzymes, the cell lysate was thermally treated at 60 ^°^C for 30 minutes before being used in the assays.

## Results and Discussion

### DE3 lysates exhibit higher PDC specific activity as compared to LysY/I^q^

We were interested in comparing the expression profiles of PDC protein from the two *E. coli* expression hosts which differ in the regulatory aspect of expression of the cloned gene under varying temperature and IPTG concentration treatments.

The overnight primary cultures of respective strain were used to start secondary cultures and IPTG was supplemented in the medium when respective culture attained OD_600_ = 0.4 (Fig 2A). At 37^°^C, the doubling time of *E. coli* is approximately 20 minutes. However, since transcription and translation are coupled in prokaryotes, the high replication rates could serve as a source of reduced protein quality due to incorrect folding of the polypeptide (de Groot & Ventura, 2006) and result in accumulation of inactive protein in the cytoplasm and/or periplasmic space (de Groot & Ventura, 2006; İncir & Kaplan, 2024; Rosano & Ceccarelli, 2014). To reduce significant accumulation of inactive protein fraction we induced the strains at 37^°^C for 2 h. It was observed that both DE3 (Fig 2B) and LysY/I^q^ (Fig 2D) strains expressed high levels of PDC at all tested concentrations of IPTG at 37 ^°^C. It seemed that level of PDC expression was relatively higher for DE3 strain as compared to LysY/I^q^. We also tested the expression profile of PDC at a reduced temperature of 18^°^C for 16 h. When strains were induced at 18^°^C, DE3 (Fig 2C) again displayed a relatively more robust expression profile as compared to LysY/I^q^ (Fig 2E). Additionally, a gradient response of protein expression directly proportional to IPTG concentration was observed in both strains at the reduced induction temperature.

**Figure 2:**
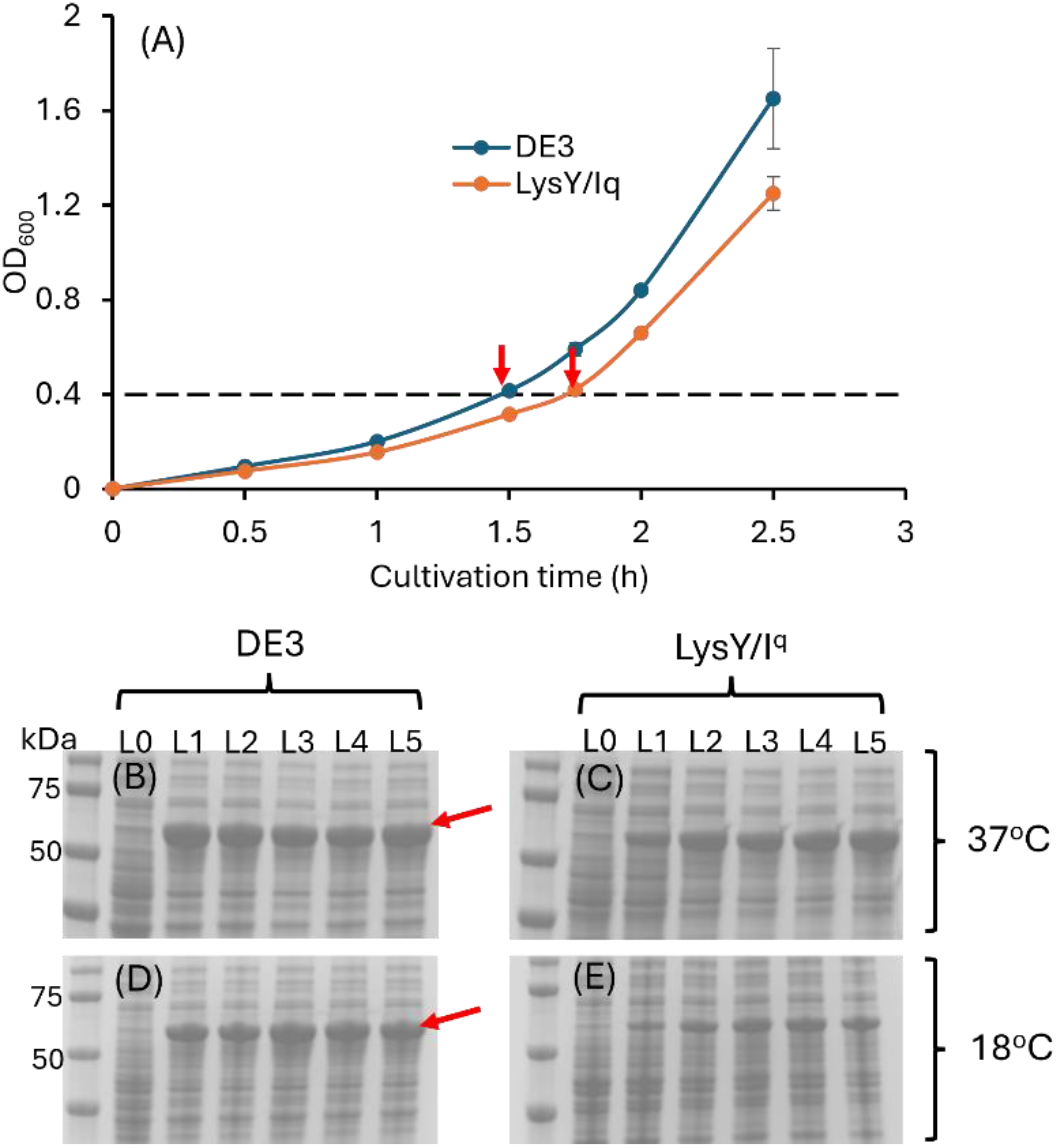
Expression of PDC in *E. coli BL21* strain(s) *DE3* and *lysY/I*^*q*^. Concentration of IPTG in lane L0, L1, L2, L3, L4 and L5 were 0, 0.1, 0.3, 0.6, 1.2 and 2.4 mM, respectively. Both strains were cultured at 37^°^C until OD_600_ reached 0.4 (A) before IPTG was added to the culture. Protein induction was performed at 37^°^C for 2 hours for *DE3* (B) and *lysY/I*^*q*^ (C). While at 18^°^C induction was performed for 16hours for *DE3* (D) and *lysY/I*^*q*^ (E). Error bars represent ± 1 S.D with average of two biological replicates.

To get an estimate of protein activity under different IPTG and temperature treatments, we performed specific activity analysis of the crude PDC lysates under each treatment. Both strains displayed distinct specific activity patterns at different temperatures. At 37^°^C, there was 0.6-fold increase in the specific activity of PDC expressed in LysY/I^q^ host when IPTG concentration was increased from 0.1 to 0.3 mM (Fig 3A). Further increase in IPTG concentrations did not result in appreciable changes in the specific activity values. Conversely, in case of strain DE3 specific activity reduced by 1.9-fold when IPTG concentration was increased from 0.1 to 0.6 mM and values did not change remarkably at further increasing concentrations of IPTG. At 18^°^C, there was remarkable increase in specific activity values of respective strain at corresponding concentration of IPTG observed at 37^°^C. The maximum specific activity in the DE3 lysates at 18^°^C was 7.8-fold higher than that observed at 37^°^C; and in case of LysY/I^q^ it was 3.1-fold higher (Fig 3B). Overall, these observations suggest three phenomena. First, at the optimum cultivation temperature of 37^°^C, increasing the expression levels of PDC in a prophage harboring strain DE3 is detrimental to its activity while in a stringently regulated strain LysY/I^q^ a moderate increase in PDC expression levels are beneficial in increasing the specific activity of the same. Second, at a reduced cultivation temperature, PDC protein constitutes a major fraction of the total protein expressed in both strains. Third, in strain DE3 the proportion of PDC out of total fraction is remarkably higher as compared to strain LysY/I^q^. To verify our observations, we proceeded to evaluate the yield and activity of purified PDC protein under different expression hosts, temperature and IPTG treatments.

**Figure 3:**
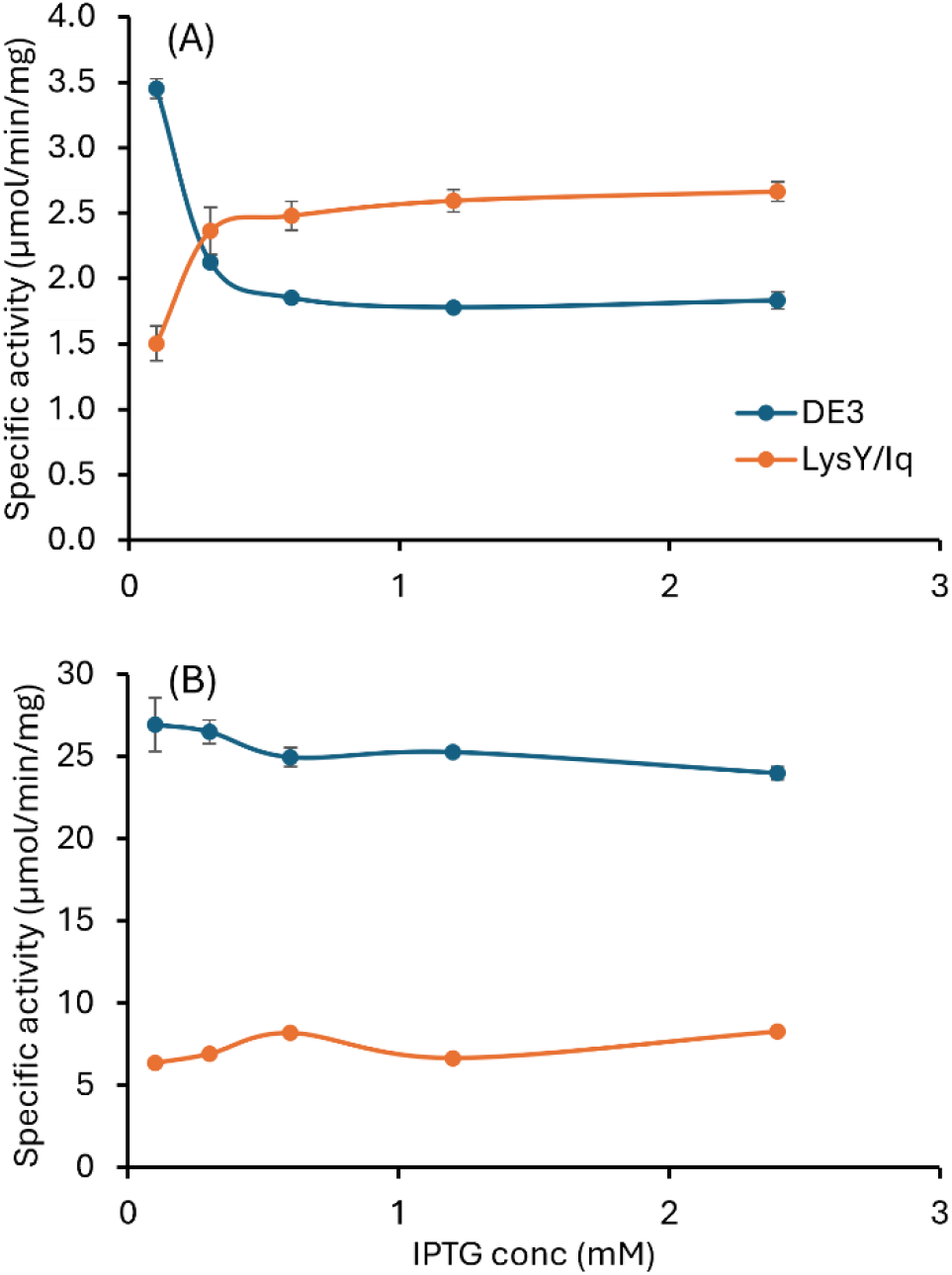
Specific activity of cell lysates from cultures cultivated at different temperatures and 0, 0.1, 0.3, 0.6, 1.2 and 2.4 mM IPTG concentrations. Cultivation was performed at either 37^°^C for two hours (A) or at 18^°^C for 16 hours (B). Equal amount of total protein was used in the assays. Results are average of two biological replicates with ±1S.D.

### Comparable PDC yield is obtained from both LysY/I^q^ and DE3

We next compared the yield of purified PDC from both strains under different temperature and IPTG treatments. To maximise the yield of purified protein we initially optimized the lysis efficiency of the cell pellet by varying the sonication time at 0.5, 1 and 2 minutes (Supplementary Fig 1). After 1 minute sonication, no PDC could be detected in the cell debris thus we selected 1 minute sonication time for further experiments. The protein purified as a single band in all purifications (Fig 4 A,B). We did not observe any evidence of leaky expression in absence of IPTG in either of the strains.

**Figure 4.**
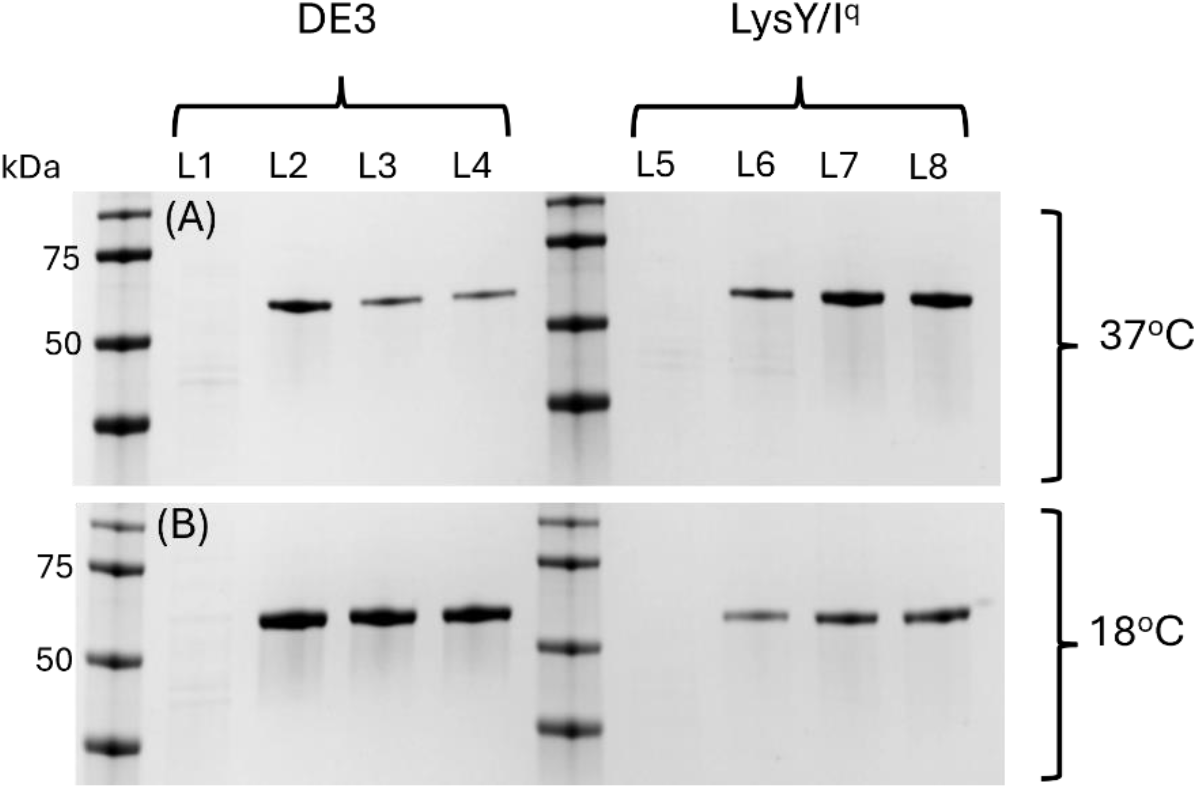
Elution profile of purified PDC protein. Protein was purified from both strains *DE3* (L1, L2, L3, L4) and *lysY/I*^*q*^ (L5, L6, L7, L8) at either 37^°^C after 2 hour induction (A) or at 18^°^C after 16 hour induction (B). Concentration of IPTG in L1, L2, L3, L4 and L5, L6, L7, L8 was 0, 0.1, 0.6, 2.4 mM, respectively. Equal volume of eluant was loaded in each well.

Upon comparing the specific activity measurements of the purified PDC from different treatments, a similar pattern of variation in specific activity values was observed as in cell lysate fractions. Expectedly, we observed an increase in the specific activity values of purified proteins as compared to cell lysate measurements. Interestingly we observed that the increase in specific activity at 37^°^C treatment for both strains (Figure 5A) was in the range of 21 to 32-fold increase as compared to 3 to 9-fold increase at 18^°^C (Figure 5B). Even though we treated the cell lysates at 60^°^C for 30 minutes for inactivation of non-specific enzymes, the dramatic increase in specific activities values suggests that at least some component of microbial cell was detrimental to specific activity measurements and was absent at a reduced cultivation temperature of 18^°^C.

**Figure 5:**
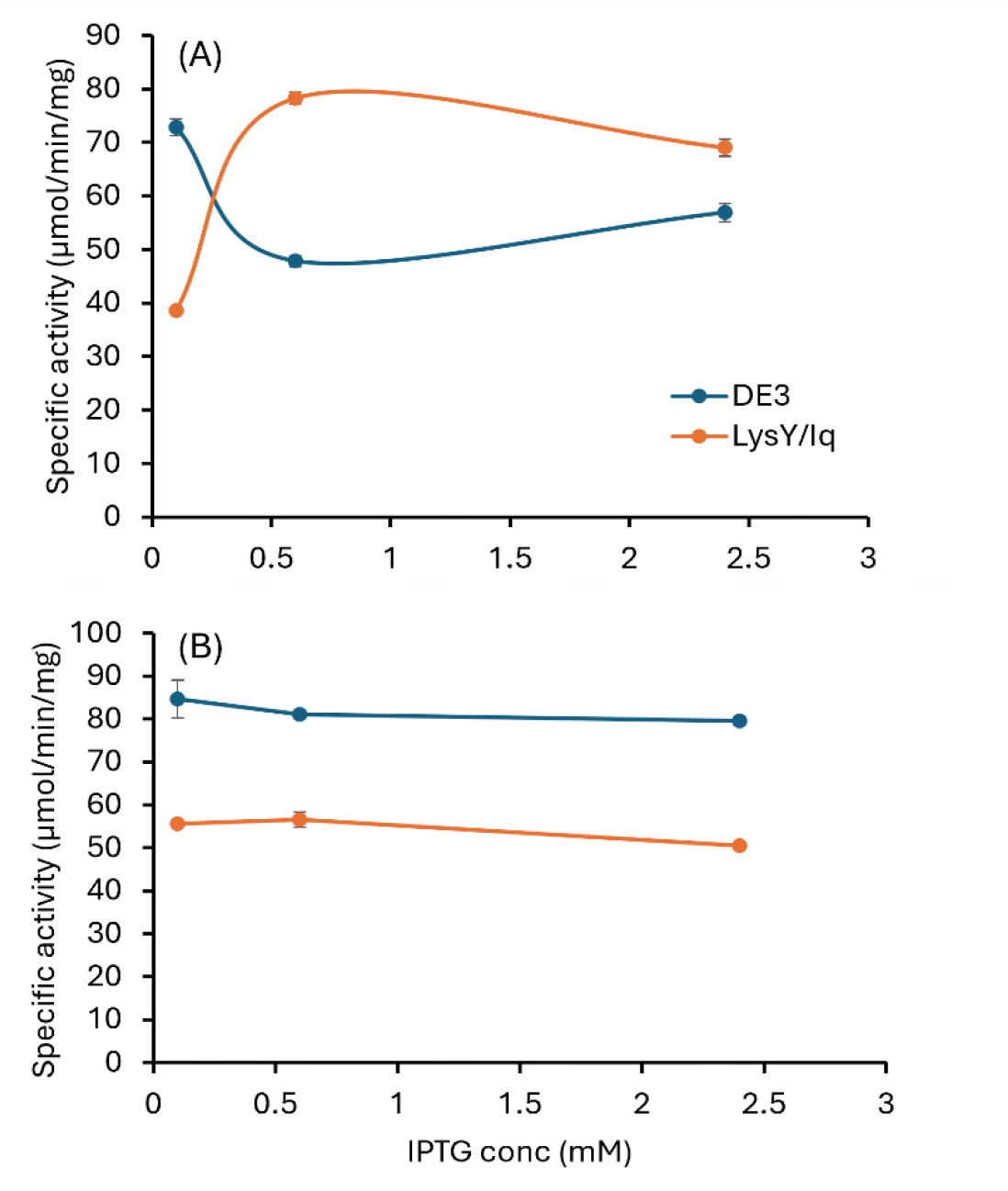
Specific activity of purified PDC protein from cultures cultivated at different temperatures and 0.1, 0.6 and 2.4 mM IPTG concentrations. Cultivation was performed at either 37^°^C for two hours (A) or at 18^°^C for 16 hours (B). Equal amount of total protein was used in the assays. Results are average of two biological replicates with ±1 S.D.

We next compared the yield of purified PDC under different cultivation treatments (TABLE 1). At 37^°^C, for strain DE3, a progressive decrease in yield of purified PDC was observed with an increase in IPTG concentration. The yield at 0.1 and 2.4 mM IPTG concentration was 0.24 (±0.03) and 0.10 (±0.02) mg of protein, respectively. It represents a decrease in yield by 58%. However, we did not observe any such remarkable variation in PDC yield for strain LysY/I^q^. For yield at 18^°^C cultivation of either strain, we did not observe any remarkable difference in the yield. A relatively high variation in the yield of the protein was observed at increasing concentration of IPTG at the lower cultivation temperature.

**TABLE 1:**
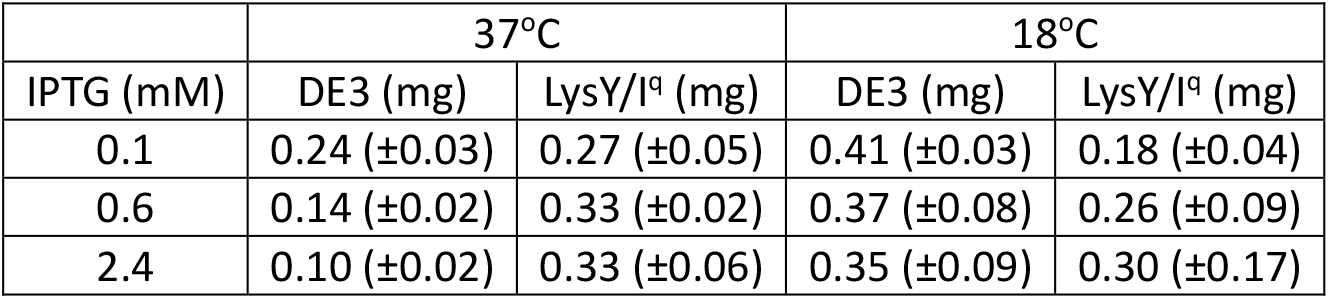
Yield of purified PDC protein obtained under different cultivation conditions. Values are average of two independent replicates with ±1S.D.

|  | 37°C |  | 18°C |  |
| --- | --- | --- | --- | --- |
| IPTG (mM) | DE3 (mg) | LysY/I<sup>q</sup> (mg) | DE3 (mg) | LysY/I<sup>q</sup> (mg) |
| 0.1 | 0.24 ($\pm 0.03$) | 0.27 ($\pm 0.05$) | 0.41 ($\pm 0.03$) | 0.18 ($\pm 0.04$) |
| 0.6 | 0.14 ($\pm 0.02$) | 0.33 ($\pm 0.02$) | 0.37 ($\pm 0.08$) | 0.26 ($\pm 0.09$) |
| 2.4 | 0.10 ($\pm 0.02$) | 0.33 ($\pm 0.06$) | 0.35 ($\pm 0.09$) | 0.30 ($\pm 0.17$) |

**TABLE 2.** Kinetic parameters of PDC purified under different treatments. Results are average of two biological replicates with ± 1 S.D.

| Temp<br>(°C) | Strain | IPTG<br>conc<br>(mM) | App. $K_m$<br>(mM) | App. $V_{max}$<br>($\text{min}^{-1}$) | $K_{cat}$<br>($\text{min}^{-1}$) | $K_{cat}/K_m$<br>($\text{mM}^{-1} \text{min}^{-1}$) |
| --- | --- | --- | --- | --- | --- | --- |
| 18°C | DE3 | 0.1 | 1.48<br>($\pm 0.11$) | 0.05<br>($\pm 0.00$) | 12,800<br>($\pm 400$) | 8,700<br>($\pm 900$) |
| | | 0.6 | 1.59<br>($\pm 0.24$) | 0.10<br>($\pm 0.02$) | 23,900<br>($\pm 4,900$) | 14,900<br>($\pm 800$) |
| | | 2.4 | 1.21<br>($\pm 0.04$) | 0.10<br>($\pm 0.01$) | 23,200<br>($\pm 3,100$) | 19,300<br>($\pm 3,200$) |
| | LysY/I<sup>q</sup> | 0.1 | 1.37<br>($\pm 0.29$) | 0.09<br>($\pm 0.01$) | 21,800<br>($\pm 2,400$) | 16,100<br>($\pm 1,700$) |
| | | 0.6 | 1.51<br>($\pm 0.08$) | 0.11<br>($\pm 0.00$) | 25,700<br>($\pm 1,200$) | 17,000<br>($\pm 100$) |
| | | 2.4 | 1.36<br>($\pm 0.24$) | 0.09<br>($\pm 0.02$) | 20,800<br>($\pm 4,300$) | 15,200<br>($\pm 500$) |
| 37°C | DE3 | 0.1 | 1.35<br>($\pm 0.07$) | 0.08<br>($\pm 0.01$) | 19,700<br>($\pm 2,100$) | 14,500<br>($\pm 800$) |
| | | 0.6 | 1.19<br>($\pm 0.06$) | 0.09<br>($\pm 0.00$) | 21,600<br>($\pm 400$) | 18,100<br>($\pm 600$) |
| | | 2.4 | 1.63<br>($\pm 0.23$) | 0.12<br>($\pm 0.02$) | 29,500<br>($\pm 4,800$) | 18,100<br>($\pm 400$) |
| | LysY/I<sup>q</sup> | 0.1 | 1.56<br>($\pm 0.39$) | 0.10<br>($\pm 0.02$) | 24,600<br>($\pm 4,700$) | 15,900<br>($\pm 1,000$) |
| | | 0.6 | 1.41<br>($\pm 0.22$) | 0.10<br>($\pm 0.01$) | 23,600<br>($\pm 3,100$) | 16,800<br>($\pm 500$) |
| | | 2.4 | 1.66<br>($\pm 0.09$) | 0.11<br>($\pm 0.01$) | 25,956<br>($\pm 1,797$) | 15,662<br>($\pm 1,921$) |

Finally, we measured the kinetic parameters of the PDC enzyme purified from the two strains. We noticed no remarkable changes in the affinity of the enzyme against its substrate pyruvate as evident by similar *apparent K*_*m*_ values determined from different purifications (Hoppner & Doelle, 1983; Raj et al., 2002). In terms of kinetic parameters we also observed that there was no remarkable difference in the values of apparent *V*_*max*_, *K*_*cat*_ and *K*_*cat*_*/K*_*m*_ ratios which suggest that the cultivation conditions do not remarkably influence the kinetic parameters of the functional PDC protein. The only exception was observed for the protein purified from strain DE3 with 0.1 mM IPTG at 18^°^C. The values of maximum reaction velocity (*K*_*m*_), turnover number of the substrate (*K*_*cat*_) and *K*_*cat*_*/K*_*m*_ ratio were approximately half of the values obtained under other purification conditions. Coincidently, it was also the treatment where the highest yield of the protein at 0.41 (±0.03) mg was reported (TABLE 1). Together, these observations suggest that the high rate of protein synthesis affected its functional properties. Since prokaryotic host has coupled transcription and translation machinery, it is likely that the balance between both processes was lost, and high yield came at a cost of functional activity. It is commonly accepted that the rate of polypeptide folding is controlled by the rate of mRNA translation (Goyal & Chaudhuri, 2015; İncir & Kaplan, 2024; Liu et al., 2022; Niwa et al., 2012). It is possible that the highest yield of PDC was achieved at the expense of improper folding of the polypeptide chain. Since at the higher IPTG concentrations at 18 ^°^C, PDC purified from strain DE3 did not display any remarkable reduction in the kinetic parameters we hypothesized that the cell responded by down modulating the protein translational machinery.

## Conclusion

. In the present study we present proof of concept that a host optimized for protein expression and harboring a tightly regulated translational mechanism can be used both quickly and efficiently to express and purify a non-toxic protein with important industrial relevance. Strain DE3 is widely used to express and purify non-toxic proteins in high yields while strain LysY/I^q^ is traditionally used for toxic proteins which are expressed and purified in relatively low yields. PDC from *Z. mobilis* is a well characterized industrial protein. In reported publications, PDC has been purified after extended cultivation hours > 10 h with not significant clarity on the yield of purified protein. We present evidence that comparable yield and kinetics of PDC can be obtained for two hour and 16 h induction time periods. Overall we report that the affinity of PDC for pyruvate (apparent *K*_*m*_), catalytic efficiency as a measured by the velocity of the reaction (apparent *V*_*max*_), turnover of the substrate (*K*_*cat*_) as well as activity at low substrate concentration (*K*_*cat*_*/K*_*m*_) are not appreciably affected by the use of different expression host, cultivation time and temperature. These results have promising applications in the cell free biology investigations where multiple purified proteins are required to scale up the complexity of the metabolic pathways. The rapid expression of heterologous proteins will significantly reduce the overall experimental turnaround time of the cell free investigations where an experimental setup with rapidly expressed (2 h) and purified proteins can be setup within a single day.Click or tap here to enter text..

## Acknowledgements

We acknowledge kind critical comments provided by Lee R. Lynd on the manuscript. Funding for this work was provided by the U.S. Department of Energy, Office of Science, Office of Biological and Environmental Research, Genomic Science Program under Award Number DE-SC0022175. This research was supported by the Center for Bioenergy Innovation (CBI), U.S. Department of Energy, Office of Science, Biological and Environmental Research Program under Award Number ERKP886.

## Author contribution

SBJ designed experiments, performed experiments, analyzed data, wrote the manuscript; DGO provided critical inputs, reviewed the manuscript, project administration and provided funding. All authors read and approved the final manuscript.

