## Supplementary data for "Rapid expression of pyruvate decarboxylase from *Zymomonas mobilis* in *E. coli* BL21 LysY/I^q^"

**Supplementary files:**

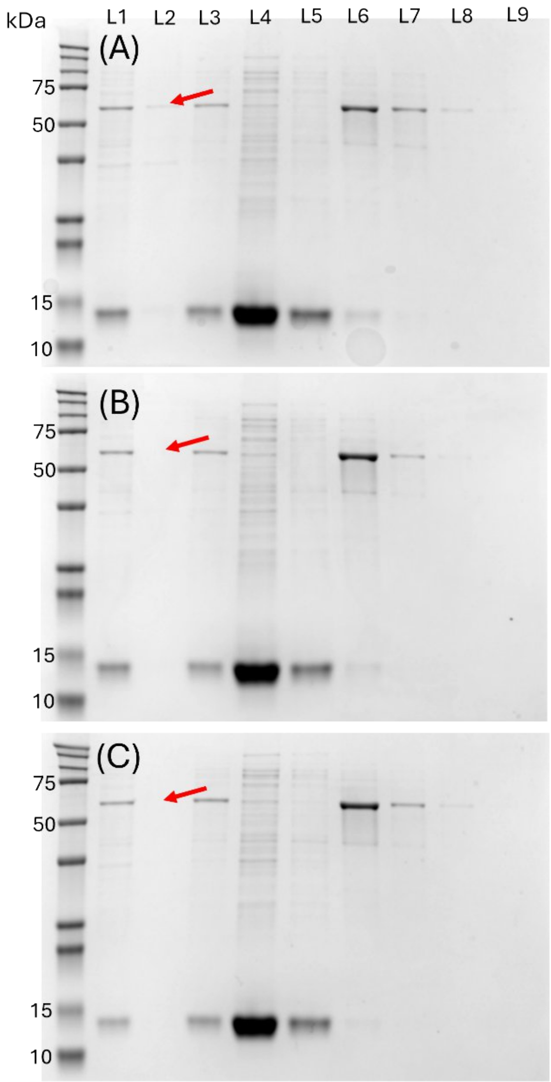


**Supplementary Figure 1. Optimizing lysis efficiency of *E. coli BL21 DE3* by sonicating cell pellets for either 30 seconds (A), 1 minute (B) and 2 minutes (C). Independent microbial cultures were induced for PDC production in presence of 0.3 mM IPTG at 18°C for 16 hours. L1 – cell lysate after sonication; L2 – cell pellet after centrifugation of cell lysate; L3 – supernatant after centrifugation of cell lysate; L4 – flowthrough after passing through column; L5 – wash buffer after passing through protein bound resin; L6, L7, L8 and L9 – elution fractions.**
